# Cross-Platform Assessment of Sub-50 nm Nanopipette Emitters for Native Electrospray Ionization Mass Spectrometry

**DOI:** 10.64898/2026.05.20.726677

**Authors:** Emily J. Byrd, Eileen J. Olivares, Zoe J. Heidersbach, Raya Sadighi, Khaled Madhoun, Iris Ansah, Matthew Kensil, Luchen Wuyang, Rafael D. Melani, Paolo Actis, Rachel R. Ogorzalek Loo, Frank Sobott, Kevin Freedman, Pei Su, Anouk M. Rijs, Antonio N. Calabrese, Joseph A. Loo

## Abstract

Native mass spectrometry (nMS) is well established for measuring protein masses and stoichiometries using nano-electrospray ionization (nESI), yet salt adduction and source activation energies can limit routine measurements. In this study, we benchmark submicron quartz nanopipette nESI emitters (<50 nm internal diameter) across three mass spectrometry platforms (quadrupole-time-of-flight, quadrupole-Orbitrap, and tribrid-Orbitrap platforms) and a wide protein range (14.5-800 kDa). We analysed the intrinsically disordered protein alpha synuclein (αS; 14.5 kDa) and holo-myoglobin (17 kDa) over a range of concentrations (10 μM-1 nM) and capillary voltages to determine limits of detection and define a gentle operating regime. We additionally observed reduced Na⁺ adduction and preservation of the Zn²⁺-bound metalloproteoform of carbonic anhydrase II (29 kDa). Proteins and protein complexes spanning the mid-to-high mass range including ovalbumin (∼44 kDa), malate dehydrogenase (∼70 kDa), transferrin (80 kDa), glutamate dehydrogenase (∼350 kDa), β-galactosidase (∼465 kDa), and GroEL (∼800 kDa), were readily detected using nanopipette emitters. Compared with conventional 1-2 μm internal diameter borosilicate emitters, quartz nanopipettes provided higher signal-to-noise ratios and fewer adducts. Finally, direct analysis of clarified bacterial lysate expressing α-synuclein yielded a clear monomeric charge-state distribution, demonstrating compatibility with complex biological matrices. Collectively, these results establish quartz nanopipette nESI as an instrument-portable, salt-tolerant approach suitable for routine nMS analysis across a broad range of protein molecular weights and sample complexities.

## Introduction

Native mass spectrometry (nMS) has become a central tool for characterising intact proteins and biomolecular assemblies by providing detailed information on mass, stoichiometry, and noncovalent ligand interactions^1–5^. Despite the broad application of nMS, two practical barriers still limit routine measurements across protein classes and instrument platforms: (i) salt adduction, which broadens peaks, obscures charge-state distributions, and impairs deconvolution, and (ii) source activation conditions required for desolvation that often trade off against spectral quality.

nMS requires gentle transfer of biomolecules such as proteins, peptides and nucleic acids from their aqueous phase into their gaseous ionic phase under non-denaturing conditions to preserve structural, conformational and non-ionic features for measurement^6–9^. This process requires the analysis of biomolecules in volatile buffers such as ammonium acetate, which does not adequately represent physiological conditions as it lacks physiological electrolytes (such as Na^+^, K^+^, PO_4_^3-^)^10–13^ that are known to be essential for maintaining the structural integrity of proteins^14,15^. Adduction of non-volatile salts and ionic species create numerous, generally nonspecific adduct peaks, which spreads the overall signal of a species over multiple peaks^12,16,17^. This results in crowding of native mass spectra with overlapping peaks, which can confound charge state distributions, reducing resolution and overall signal-to-noise. These issues are amplified for higher-mass assemblies and in more complex matrices, which has motivated the exploration of strategies to improve desolvation, whilst preserving weak, noncovalent interactions^18–20^.

Additionally, the presence of salts and non-volatile species in solutions being analysed by nMS can generate salt clusters in the lower *m/z* range of mass spectra, preventing detection of higher charge states or smaller analytes and can prevent the observation of analytes altogether due to ion suppression^10,21^. Therefore, non-volatile buffer components typically need to be removed from analyte samples prior to nMS analyses. This is usually achieved by dialysis methods or on-column desalting/buffer exchange procedures, which can ultimately lead to considerable sample loss, aggregation or subsequent precipitation of proteins and loss of structural features, protein-protein assemblies and protein-ligand complexes.

Submicron-sized nanoelectrospray ionization (nESI) emitters have been applied for nMS analysis of proteins from biochemical buffers^22–32^ and in the presence of detergents^33^ in attempts to circumvent the need for laborious buffer exchange protocols. Alternative desalting approaches include the implementation of dual-barrel nanoemitters to direct nMS of proteins in biochemical buffers through incomplete mixing from a single Taylor cone^34–36^. In this set-up, mixing times can be controlled by varying tip diameter and geometry and results in the generation of salt-depleted droplets with smaller radii than conventional nESI, preserving noncovalent assemblies^37^. Online buffer exchange is an alternative approach that relies on chromatographic separation and dilution into a nMS compatible running buffer (ammonium acetate)^38–41^. However, this approach does not enable analyses in biochemical buffer systems and generates a short time window for spectra acquisition during analyte elution windows.

Submicron-sized nESI emitters using quartz nanopipettes also directly address these strategies. Operating at low flow rates from quartz tips <50 nm inner diameter (I.D.), nanopipettes generate nanodroplets with reduced droplet radii compared to standard borosilicate emitters, providing on-tip desalting and improved signal-to-noise without offline cleanup^24,31,32,42^. We have previously demonstrated the detection of resolvable charge state distributions arising from the intrinsically disordered protein α-synuclein directly from biochemical buffer systems^43^.

Here we deliver a cross-platform benchmark of submicron quartz nanopipette nESI using a panel of proteins spanning diverse molecular weights (MWs) and including higher-order assemblies; namely, alpha-synuclein (αS, ∼14.5 kDa), holo-myoglobin (∼17 kDa), carbonic anhydrase II (∼29 kDa), ovalbumin (∼44 kDa), malate dehydrogenase dimer (∼70 kDa), transferrin monomer (80 kDa), glutamate dehydrogenase hexamer (∼335 kDa), β-galactosidase tetramer (∼465 kDa) and GroEL tetradecamer (∼800 kDa). We evaluate performance on four instrument families: Waters Q-TOF Synapt G2-Si, ThermoFisher Scientific Q-Exactive Orbitrap UHMR, Orbitrap Ascend and Bruker timsTOF Pro 2, and two conventional ways of electrical contact: gold/titanium coated nESI emitters and platinum wire charging of the analyte solution^32,42^. These results demonstrate that nanopipettes are a versatile nESI modality with stable performance, agnostic of vendor-specific ESI interface design. By comparison of quartz nanopipettes to conventional 1–2 μm borosilicate emitters, including analysis of a range of capillary voltages, we map a “gentle” operating window on each platform. These comparisons address a long-standing concern that ultranarrow tips might perturb native structures and establish when submicron emitters provide clear advantages over standard tips for different mass regimes. Also, we show that recombinant proteins can be detected from crude bacterial cell lysates for protein identification.

Together, this study provides a multi-instrument assessment of quartz nanopipette nESI across a broad protein mass range, evidence for robust on-tip desalting and improved native spectral quality, guidelines for emitter operating conditions by protein class and platform and an application vignette demonstrating proteoform-level analysis directly from crude bacterial lysates. By standardizing amenable analytes and operating conditions, we aim to position submicron quartz nanopipettes as a practical, portable front end for high-quality native MS across laboratories and instruments.

## Methods

### Sample preparation

Myoglobin (SKU: M0630; equine skeletal muscle), carbonic anhydrase II (CAH II - SKU: C2624); bovine erythrocytes), ovalbumin (SKU: A-2513; Grade VI - chicken egg), L-glutamate dehydrogenase (GDH - SKU: G7882; bovine liver), β-galactosidase (β-Gal - SKU: G5635; E. Coli), Chaperonin 60 (GroEL - SKU: C7688; E. Coli), and L-malate dehydrogenase (LMDH - SKU: 442610-10KU; pig heart - mitochondrial) were purchased from Sigma-Aldrich. Myoglobin and CAH II were dissolved in 100 mM ammonium acetate (pH 6.6-6.8) and analysed without buffer exchanging. Ovalbumin, GDH, β-Gal and GroEL were prepared for mass spectrometry by dissolving each protein in 200mM ammonium acetate (pH 6.6-6.8). Prior to analysis, ovalbumin, GDH, β-Gal and GroEL were buffer exchanged into 200mM ammonium acetate using appropriate Amicon® Ultra centrifugal filters with the appropriate molecular weight cut off (MWCO). Filters were conditioned before use, and buffer exchange was performed two to four times. Samples were centrifuged at 10,000 x g for five to ten minutes per exchange cycle. Holo transferrin (Holo Tf - SKU: 616397; Human plasma, Sigma Aldrich) and apo transferrin (Apo Tf - Cat. #: 16-16-A32001 Human plasma, Athens Bioscience) were buffer exchanged using the same procedure as above and prepared in 500 mM ammonium acetate. αS (AB-600-100, wild type human, Recombinant, Alexo-Tech) and was prepared in 50 mM ammonium acetate, without prior buffer exchange. Proteins were analyzed at concentrations of 10 µM or lower where stated.

### Preparation of bacterial cell lysates

Competent BL21-DE3 *E. coli* cells expressing NatB for N-terminal acetylation were generated as descried previously^44,45^. Competent NatB-BL21 DE3 cells were transformed with a pET-23a plasmid encoding wild type human full length α-synuclein (αS) to express both NatB and αS for N-terminal acetylation (αSNTA) or were transformed with a pET-28a vehicle control. Overnight cell cultures were grown in 10 mL LB media supplemented with 25 µg/mL chloramphenicol and 100 µg/mL carbenicillin. 0.5 mL of the overnight culture was used to inoculate 50 mL of LB media with 25 µg/mL chloramphenicol and 100 µg/mL carbenicillin until an OD_600_ of 0.6 was reached and protein expression was induced with the addition of 0.01 mg/mL Isopropyl β-D-1-thiogalactopyranoside (IPTG) for 3 hr at 37°C, 200 rpm. The cultures were centrifuged at 4,000 x g for 15 min. Cell pellets were washed once with 100 mM ammonium acetate (pH 6.6-6.8). Cells were then resuspended in 2 mL of buffer containing 100 mM ammonium acetate, pH 6.6.-6.8 and protease inhibitor cocktail (Roche) and lysed by sonication. The lysate was centrifuged at 11,000 x g for 30 min. The supernatant was stored at −80°C and diluted 1:1 with 100 mM ammonium acetate before direct nMS.

### Nanopipette nESI fabrication

Nanopipette nESI emitter tips were fabricated using 1.0 mm outer diameter and 0.5 mm inner diameter quartz capillaries (QF100-50-7.5; Sutter Instrument) with a P2000 laser puller (Sutter Instrument). A two-line protocol was used: line 1 with HEAT 750, FIL 4, VEL 30, DEL 150, PUL 80, followed by line 2 with HEAT 850, FIL 3, VEL 40, DEL 135, PUL 225. The pulling protocol is specific to the instrument and can vary between different pullers. For nMS measurements using uncoated nanopipettes, the emitters were loaded with analyte solution (3-5 µL) and fitted with a platinum wire (PT00-WR-000117; Goodfellow) prior to use. Figure S1a shows a representative SEM micrograph of the nanopipettes used in this study.

Coated nanopipette nESI emitter tips (Figure S2) used for transferrin analyses were fabricated using 1.0 mm outer diameter and 0.7 mm inner diameter quartz capillaries (QF100-70-7.5; Sutter Instrument) with a P2000 laser puller (Sutter Instrument). A one-line protocol was used: line 1 with HEAT 630, FIL 4, VEL 61, DEL 125, PUL 120. The pulling protocol is specific to the instrument and can vary between different pullers. Coated nanopipettes (∼20 nm) were prepared using a sputter coater (Desk V, Denton Vacuum, LLC) with argon as the sputtering gas. After plasma cleaning the nanopipettes for 10 minutes, a thin titanium adhesion layer was deposited onto the outer glass surface (90 seconds) followed by gold deposition for 30 seconds. 3-5 μL of protein solution was loaded into a nanopipette nESI capillary emitters.

### Standard nESI emitter fabrication

Standard nESI emitter tips were fabricated using 1.2 mm outer diameter and 0.69 mm inner diameter borosilicate glass capillaries (EC1-30-0042; Harvard Apparatus) with the P-1000 micropipette puller (Sutter). A one-line protocol was used: HEAT 478/PULL 0/ VEL 65/ DELAY 80/ PRESSURE 300. Figure S1b shows a representative SEM micrograph of the standard nESI emitters used in this study.

### Synapt G2-Si

Native MS experiments performed on a Waters Synapt G2-Si mass spectrometer with travelling (T-wave) ion mobility and a nano-ESI source. Platinum wire (PT00-WR-000117; Goodfellow, diameter 0.125mm, GoodFellow) was secured in the source housing so that the wire could be fitted into the nanopipette emitter. Instrument parameters were set as follows: capillary voltage 0.2-0.6 kV (increased up to 2.0 kV for demonstration of operating windows), source temperature 30 °C, sampling cone 18 V, extraction cone 1.0 V, trap collision energy 5 V, transfer collision energy 2.0 V, trap DC bias 30 V (10 V in the case of CAH II to preserve Zn^2+^ cofactor binding). For ion mobility measurements, IM wave velocity 300 m/s, IM wave height 7.0 V. Gas pressures in the instrument were: trap cell 0.0258 mbar, IM cell 0.36 mbar. For analysis of bacterial lysates, the crude lysates were diluted 1:1 with 100 mM ammonium acetate prior to analysis. Raw data was extracted using MassLynx V4.2 and processed manually.

### Q-Exactive Orbitrap UHMR

Ovalbumin, GDH, β-Gal and GroEL samples were analysed using a Q Exactive UHMR Orbitrap mass spectrometer (Thermo Fisher Scientific) modified for the transmission and detection of high *m*/*z* ions. All spectra were calibrated externally, using caesium iodide. 3-5 μL of protein solution was loaded into a nanopipette nESI capillary emitters. Platinum wire (PT00-WR-000117; Goodfellow, diameter 0.125mm, GoodFellow) was secured in the source housing so that the wire could be fitted into the nanopipette emitter.

The source was operated in positive mode with a capillary voltage of 0.2-0.4 kV and an inlet capillary temperature of 250°C. Mass spectra were recorded at resolutions of 6250 and 12500, with an AGC target of 1×10^6^, a max injection time of 100.0 ms, and extended trapping set to −1.0 eV. The source offset was set to 21.0 V. In-source trapping was enabled with a desolvation voltage ranging from −10V to −200V. Ion transfer m/z optimization parameters were as follows: ion transfer target set to high m/z; inj. fl. RF ampl., 700 V; bent. fl. RF ampl., 940 V; trans. MP and HCD-cell RF ampl, 900V; C-trap RF ampl., 2950 V; injection flatapole DC, 5.0 V; inter flatapole lens, 4.0 V; bent flatapole DC, 2.0 V; transfer multipole lens, 0V; and C-trap entrance lens, 2 V.HCD events configured with a purge time of 5 ms, a C-trap exit lens voltge of 2 V, an HCD field gradient of 70.0 V, and a trapping gas pressure of 5.0-7.0 V. Raw data was extracted from FreeStyle 1.8 (ThermoFisher Sceintific) and analysed manually.

### Orbitrap Ascend

MDH was analysed using a hybrid quadrupole–linear ion trap–Orbitrap mass spectrometer (Orbitrap Ascend Structural Biology Tribrid; Thermo Fisher Scientific). 3-5 μL of protein solution was loaded into a nanopipette nESI capillary emitters. Platinum wire (PT00-WR-000117; Goodfellow, diameter 0.125mm, GoodFellow) was secured in the source housing so that the wire could be fitted into the nanopipette emitter.

The source was operated in positive mode with a capillary voltage of 0.4-1.4 kV and an inlet capillary temperature of 275°C. Mass spectra were recorded at a resolution of 30,000, with an AGC target of 5×10^5^ and an ion injection time of 527.123 ms. In-source trapping was enabled with a desolvation voltage ranging from 50 V. Raw data was extracted from FreeStyle 1.8 (ThermoFisher Sceintific) and analysed manually.

Further experiments for Holo and Apo Tf were performed using a hybrid quadrupole–linear ion trap– Orbitrap mass spectrometer (Orbitrap Ascend Biopharma Tribrid; Thermo Fisher Scientific). 3-5 μL of protein solution was loaded into coated nanopipette nESI capillary emitters.

The source was operated in positive mode with a capillary voltage of 0.8-1.0 kV and an inlet capillary temperature of 275°C. Ensemble-mode mass spectra were recorded at a resolution of 90,000, with an AGC target of 5×10^5^ and an ion injection time of 200 ms. In-source CID was enabled with a voltage of 65V and a scaling factor of 0.02. Raw data was extracted from FreeStyle 1.8 (ThermoFisher Scientific) and analysed manually or using UniDec^46^. Direct Mass Technology (DMT) mode was performed using the same ion source and frontend optics settings, and data was acquired with a transient length of 0.5 s.

### timsTOF Pro 2

αS measurements were performed on a timsTOF Pro 2 mass spectrometer (TIMS-Qq-TOF, Bruker Daltonics) operated in positive-ion mode over an m/z range of 1000–5000. Monomeric αS solution (3– 5 μL) was loaded into a nanopipette nESI capillary emitter. Samples were introduced via a home-built nESI source based on a previously reported design^47^. Platinum wire (PT00-WR-000117; Goodfellow, diameter 0.125mm, GoodFellow) was secured in the source housing so that the wire could be fitted into the nanopipette emitter.

The instrument was operated at a tunnel-in pressure of 2.6 mbar and was calibrated for both mass and ion mobility using Agilent ESI Tune Mix under the corresponding tunnel-in pressure conditions. The source capillary voltage was set to 1800 V, and the source temperature was maintained at 180 °C. Ion funnel RF amplitudes were set to 300 Vpp and 600 Vpp for funnels 1 and 2, respectively, and the in-source CID voltage was set to 30 V.

For TIMS-MS measurements optimised setting for αS were used^48^. The Δ potentials were set to Δ1 = 50 V, Δ2 = −150 V, Δ3 = 70 V, Δ4 = 100 V, Δ5 = 0 V, and Δ6 = 180 V. The TIMS ramp time was 100 ms, with an accumulation time of 50 ms. Ion mobility was measured over a mobility range of 1/K₀=0.9-2.0 V.s/cm².Mass spectra and extracted ion mobilograms were processed using Bruker Compass DataAnalysis v6.1. Following charge-state assignment of the extracted ion mobility distributions, experimental TIMS CCS distributions were determined for each detected species.

### Scanning Electron Microscopy (SEM)

The pore dimensions of the standard nESI and nanopipette nESI emitters were characterised uncoated by field emission SEM with a FEI Nova 450 at an accelerating voltage of 3–5 kV. Images were prepared using a CBS detector (back-scattered electron detector).

## Results and discussion

### Low molecular weight proteins Myoglobin, CAH II and Ovalbumin

We developed protocols for the reproducible fabrication and characterization of sub-50 nm using scanning electron microscopy (SEM) and electrical measurements (Fig S1). Electron microscopy indicated that the outer diameter of the nanopipette was ∼30 nm suggesting that the nanopore diameter should be ∼20 nm (Fig S1a). SEM is a powerful tool of the direct imaging of nanopipettes, but it is well established that the electron beam causes shrinkage of the nanopore and could lead to an underestimation of its size^49^. We have therefore performed an electrical characterization of the nanopipette emitters by measuring their current-voltage (i-V) response in 0.1 M KCl (Fig S1c). This approach allows the nanopore size to be estimated by calculating its electrical resistance^50^. Assuming an inner cone angle of the nanopipette of ∼2°, we estimated a nanopore diameter of ∼20 nm, which is consistent with the SEM measurements.

We first assessed the performance of the nanopipette nESI emitters with standard lower molecular weight proteins. Using the Synapt G2-Si Q-TOF platform, nanopipettes produced consistently stable electrospray (Figure S3) at markedly lower capillary potentials (∼0.4 kV) than standard 1–2 µm emitters (∼1.0-1.4 kV), yielding native charge state distributions for holo-myoglobin and carbonic anhydrase II measured in ammonium acetate without performing any buffer exchange/desalting procedures (Figure 1). We observed a clear reduction in Na^+^ adducts to both proteins when nanopipette emitters were used.

**Figure 1.**
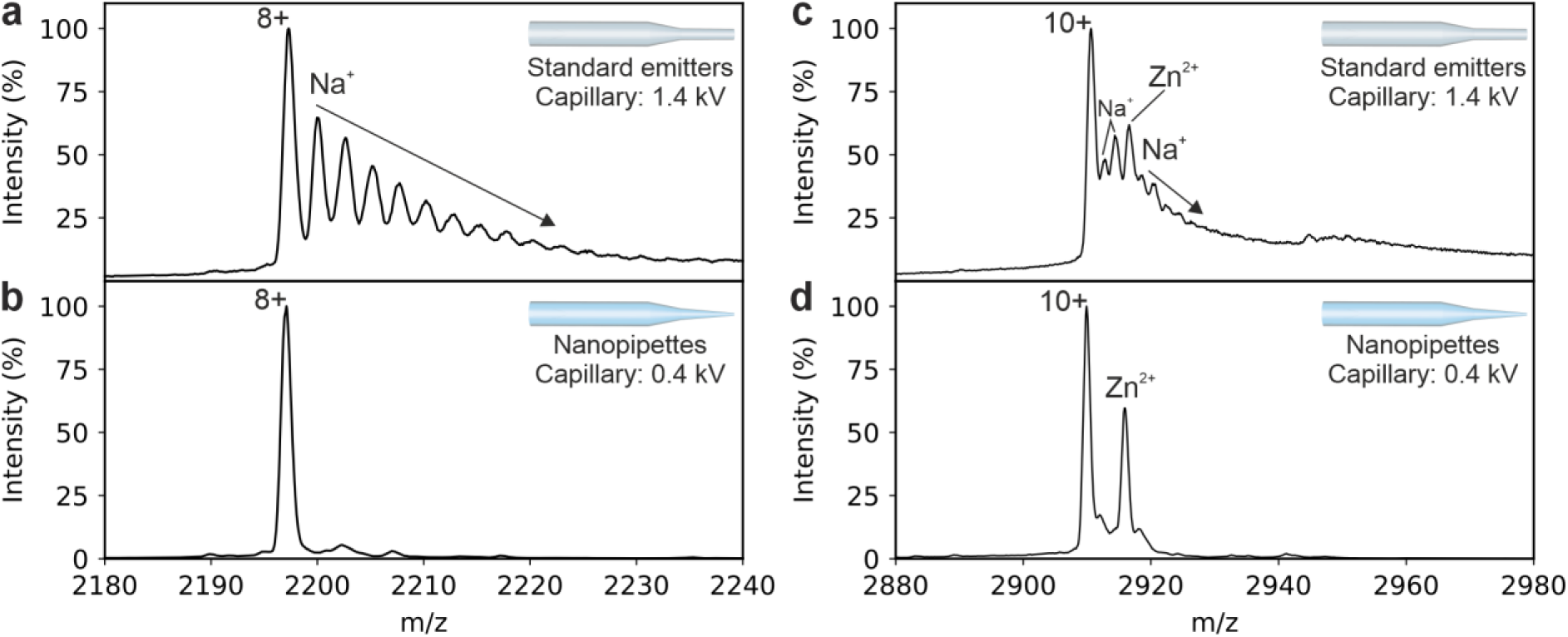
Nanopipette nESI improves native mass spectra of low-mass proteins by suppressing adducts while preserving cofactor binding. (a) Holo-myoglobin spectrum showing the 8+ charge state acquired using standard emitters (capillary 1.4 kV) with up to eight Na⁺ adducts. (b) Myoglobin 8+ charge state acquired using nanopipette emitters (capillary 0.4 kV) with markedly reduced Na⁺ adduction. (c) Carbonic anhydrase II 10+ charge state acquired using standard emitters (1.4 kV), nonspecific Na⁺ adducts overlap with the Zn²⁺ metalloproteoform. (d) Carbonic anhydrase II 10+ charge state acquired using nanopipette emitters (0.4 kV). All spectra were acquired in 100 mM ammonium acetate (pH 6.6-6.8) without prior desalting procedures on a Synapt G2-Si; Schematics (inset) show emitters used and annotations indicate operating voltages.

In the case of holo-myoglobin, the dominant 8+ charge state acquired with standard emitters (Figure 1a) showed multiple Na⁺ adduct peaks (>8 adducts). Under otherwise matched conditions except for a lower capillary voltage (0.4 kV), nanopipettes remarkably resolved the isolated, un-adducted 8+ peak, while retaining the native holo heme-bound population, consistent with gentler transfer and on-tip desalting whilst maintaining cofactor interactions (Figure 1a,b). For carbonic anhydrase II, standard emitters produced a 10+ charge state broadened by Na⁺ adduction that overlaps with the Zn²⁺ metalloproteoform (Figure 1c). With nanopipettes, nonspecific Na⁺ adducts were removed, and the Zn²⁺-bound proteoform was preserved. This selective retention of specific metal binding alongside loss of nonspecific adduction reflects the on-tip desalting potential of nanopipettes nESI emitters. Full spectra are shown in Figure S4, evidencing the improvement in signal-to-noise through the minimisation of overall baseline noise.

Additionally, we probed the effects of different capillary voltages on obtaining myoglobin signal from standard emitters and nanopipette emitters (Figure S5 and Figure S6). Nanopipettes require lower voltages (0.2-1.0 kV) to observe comparable signal intensity to standard emitters (1.0-2.0 kV).

To determine the limit of detection of holo-myoglobin, a concentration series was measured for a two-minute acquisition time. A resolvable charge state distribution for native myoglobin was measured under low capillary voltages down to 10 nM protein concentration (Figure 2). From this analysis along with the ability to generate long-lived, stable electrosprays (observed for CAH II in Figure S3), we can postulate that the quartz surface of nanopipette emitters enables the observation of low abundant proteins with minimal sample loss to adsorption^26,51–53^.

**Figure 2.**
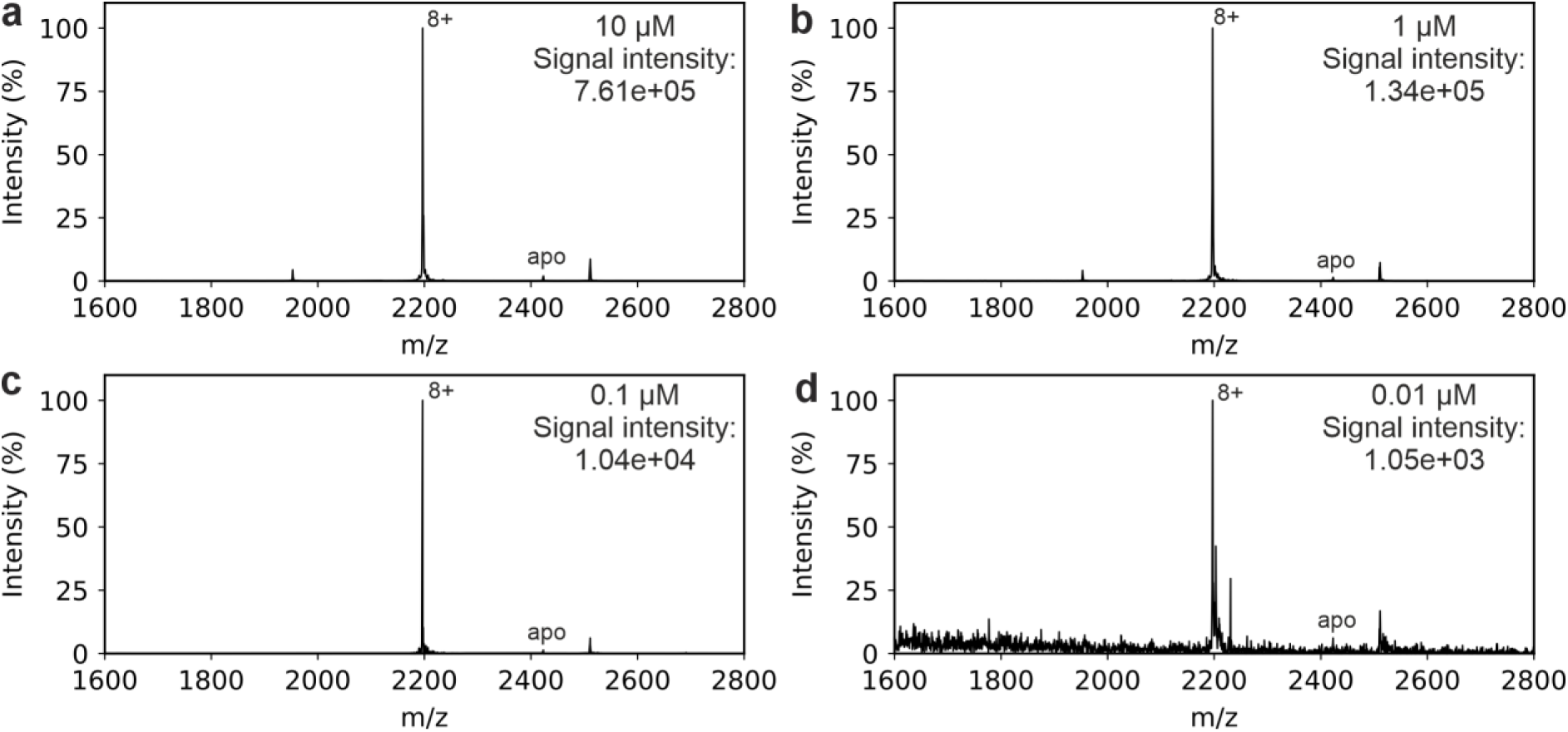
Myoglobin concentration series using nanopipette emitters. Native mass spectra of myoglobin at protein concentrations of (a) 10 μM, (b) 1 μM, (c) 0.1 μM, (d) 0.01 μM. The total ion count (TIC) from the combined two-minute spectral acquisition is denoted in the upper right-hand side of each panel. All spectra were acquired in 100 mM ammonium acetate (pH 6.6-6.8) on a Waters Synapt G2-Si.

A comparison of nMS with the nanopipette emitters on two different instrument platforms was performed using ovalbumin (∼44 kDa) in 100 mM ammonium acetate as the test subject on both the Synapt G2-Si and the Q Exactive UHMR Orbitrap (Figure 3). Analyses on both instruments revealed that nanopipettes produced a stable nanoelectrospray at low capillary voltages, delivering comparable charge state distributions, and confirming that nanopipettes can be applied across vendor platforms. The 11+ charge state displayed comparable peak distributions representing various glycoforms^54,55^ on both platforms, indicating comparable desalting efficiency and low in-source activation requirements (capillary voltage of 0.6 kV). Together, with our observations of myoglobin and CAH II, these data establish that sub-50 nm nanopipettes improve spectral quality for low-mass proteins by suppressing nonspecific salt adduction while preserving native cofactors and can be applied across platforms with improved signal-to-noise without extensive re-optimization at significantly lower capillary voltages than conventional emitters.

**Figure 3.**
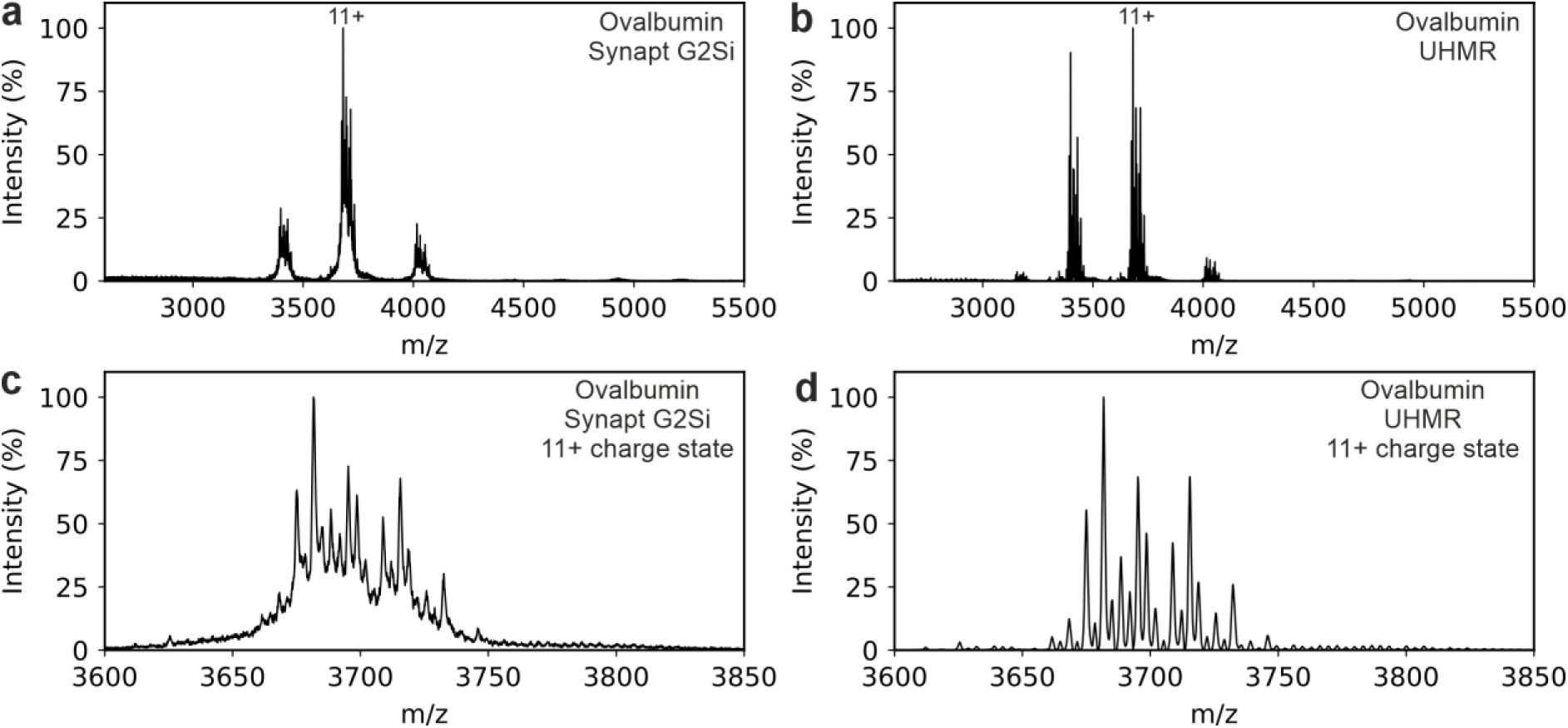
Ovalbumin native mass spectra acquired with nanopipette emitters are consistent across nMS platofrms. (a) Mass spectrum of ovalbumin acquired on a Waters Synapt G2-Si showing a resolved charge-state distribution from 10+ to 12+. (b) Mass spectrum of ovalbumin acquired on a ThermoFisher Scientific Q Exactive UHMR Orbitrap with charge states 10+ to 13+ acquired at 12,500 Orbitrap resolution. Expansion of the 11+ state from mass spectra acquired on the (c) Synapt G2-Si and (d) UHMR. All data were acquired in 200 mM ammonium acetate using <50 nm I.D. nanopipette emitters and a capillary voltage of 0.6 kV.

### Analysis of mid-mass range protein assemblies

The measurement of mid-mass, noncovalent protein assemblies were evaluated using the Orbitrap UHMR platform and the Orbitrap Ascend. To further assess nanopipette performance for low-abundance protein detection and metalloproteoform preservation, apo- and holo-transferrin were analysed using coated nanopipettes in 500 mM ammonium acetate using and Orbitrap Ascend platform. Coated tips used for these measurements had an outer diameter of 20 nm and an inner diameter of 10 nm.

At 1 µM, both apo- and holo-transferrin produced well-resolved native charge-state distributions, and deconvolution revealed the expected mass shift corresponding to binding of two Fe³⁺ ions when comparing the apo and holo forms (Figure 4a-c). Notably, apo-transferrin remained detectable at 1 nM, demonstrating the sensitivity of the coated nanopipette approach (Figure 4d). Single-scan measurements at this concentration revealed discrete individual ion events, and Direct Mass Technology (DMT) mode analysis resolved multiple transferrin-derived mass populations, including species corresponding to the N- and C-terminal lobes (∼40 kDa), intact monomeric transferrin (∼80 kDa), and dimeric transferrin (∼160 kDa; Figure 4e-f). These data highlight the compatibility of nanopipette nESI with both native metalloprotein analysis and single-ion detection workflows, extending the benchmark beyond ensemble-averaged spectra towards measurement of dilute, heterogeneous protein populations, and also demonstrate that nanopipette emitters work efficiently with gold coating without tip blocking.

**Figure 4.**
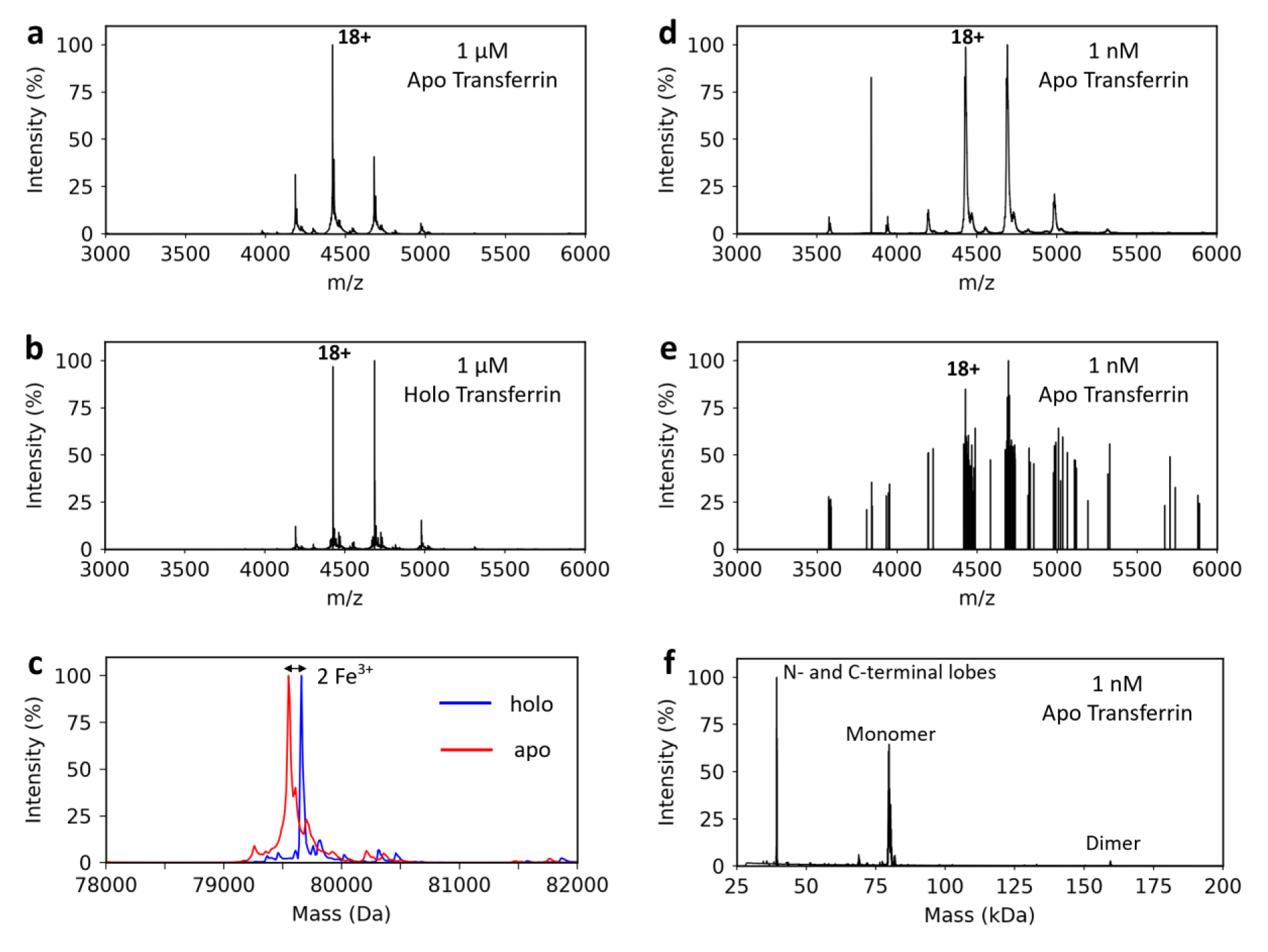
Nanopipette analysis of transferrin. (a-b) Representative averaged spectra of 1 µM apo-transferrin and 1 µM holo-transferrin, respectively, in 500 mM Ammonium Acetate solution using gold coated nanopipettes measured on a ThermoFisher Scientific Orbitrap Ascend Structural Biology Tribrid. (c) Deconvoluted spectra of a) and b) comparing apo- and holo-transferrin, highlighting the shift associated with the binding of the two ferric ions in holo-transferrin.(d) Representative averaged spectra of 1 nM apo-transferrin in 500 mM ammonium acetate solution. (e) Representative single-scan measurement demonstrating the detection of individual transferrin ion events at 1 nM. (f) DMT spectrum of the single ion detection of 1 nM apo-transferrin revealing distinct mass populations corresponding to the N- and C-terminal lobes (∼40 kDa), intact monomeric transferrin (∼80 kDa), and dimeric transferrin (∼160 kDa).

Analysis of glutamate dehydrogenase (GDH, homohexamer ∼335 kDa) in 200 mM ammonium acetate, using an Orbitrap UHMR, produced a well-resolved native charge-state distribution with no significant dissociation products (i.e. monomeric GDH), enabling assignment of the intact hexamer with a dominant charge state of 41+ (Figure 5a). Under identical instrument parameters, β-galactosidase (β-Gal, homotetramer ∼465 kDa) displayed an equally resolvable, intact charge state distribution corresponding to the intact tetramer (Figure 5b). Nanopipettes supported stable spray at lower capillary potentials (0.7-0.8 kV) than required for standard 1-2 µm emitters (Figure S7 example for β-Gal). The diameter of the GDH hexameric assembly is ∼ 8-8.6 nm^56,57^ and the diameter of the β-Gal tetramer is ∼ 18 nm; and we show here that these protein assemblies are amenable to analysis from ∼20 nm I.D. nanopipette emitters. These observations not only expand the application of nanopipette emitters across increasing protein MWs, but they also demonstrate stable electrosprays with limited blocking of the nanopipette emitters (Figure S7), which offers attractive potential of applying nanopipettes for analyses that require long acquisitions times.

**Figure 5.**
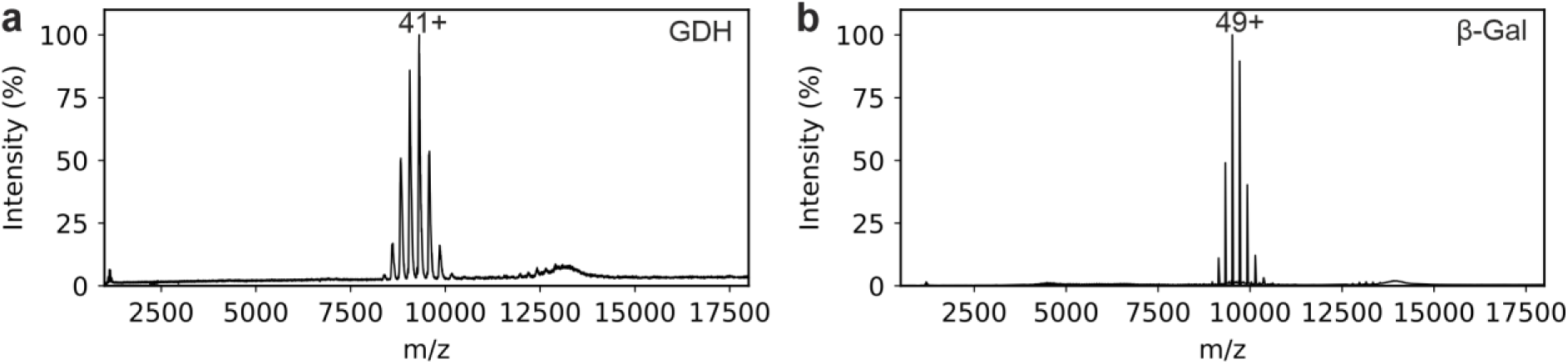
Nanopipette nESI of GDH and β-Gal protein assemblies. Mass spectra of (a) Glutamate dehydrogenase (GDH) hexamer (∼350 kDa) and (b) β-galactosidase (β-Gal) tetramer (∼465 kDa). Both spectra were acquired in 200 mM ammonium acetate buffer (pH 6.6-6.8), at 10 µM protein concentration, on a Q-Exactive Orbitrap UHMR platform.

### High-mass assembly detection: GroEL

To test the upper mass limits of nanopipette emitters, we analyzed the GroEL tetradecamer (∼800 kDa) on a ThermoFisher Scientific Q-Exactive Orbitrap UHMR. Under gentle native conditions at low capillary voltages (0.4-0.8 kV), nanopipettes produced a well-resolved charge-state distribution with a dominant charge state of 67+^58,59^ with no dissociation products, enabling straightforward charge-state assignment to a mass consistent with the intact GroEL tetradecamer. We performed a concentration series spanning 10 μM to 10 nM (Fig. 6) that showed the preserved intact GroEL tetradecamer complex and its characteristic charge distribution^58,59^. At 0.01 μM protein concentration, the GroEL charge state distribution remained consistent, indicating that low flow (the flow rate via electrospray is estimated at 36 nL min^-1^ and 18 nL min^-1^ for borosilicate emitters with I.D’s of 1.84 µm and 684 nm, respectively^60^) nanopipette nESI maintains efficient transmission and gentle desolvation even for higher MW protein assemblies at sub-micromolar sample concentrations (Figure S8).

**Figure 6.**
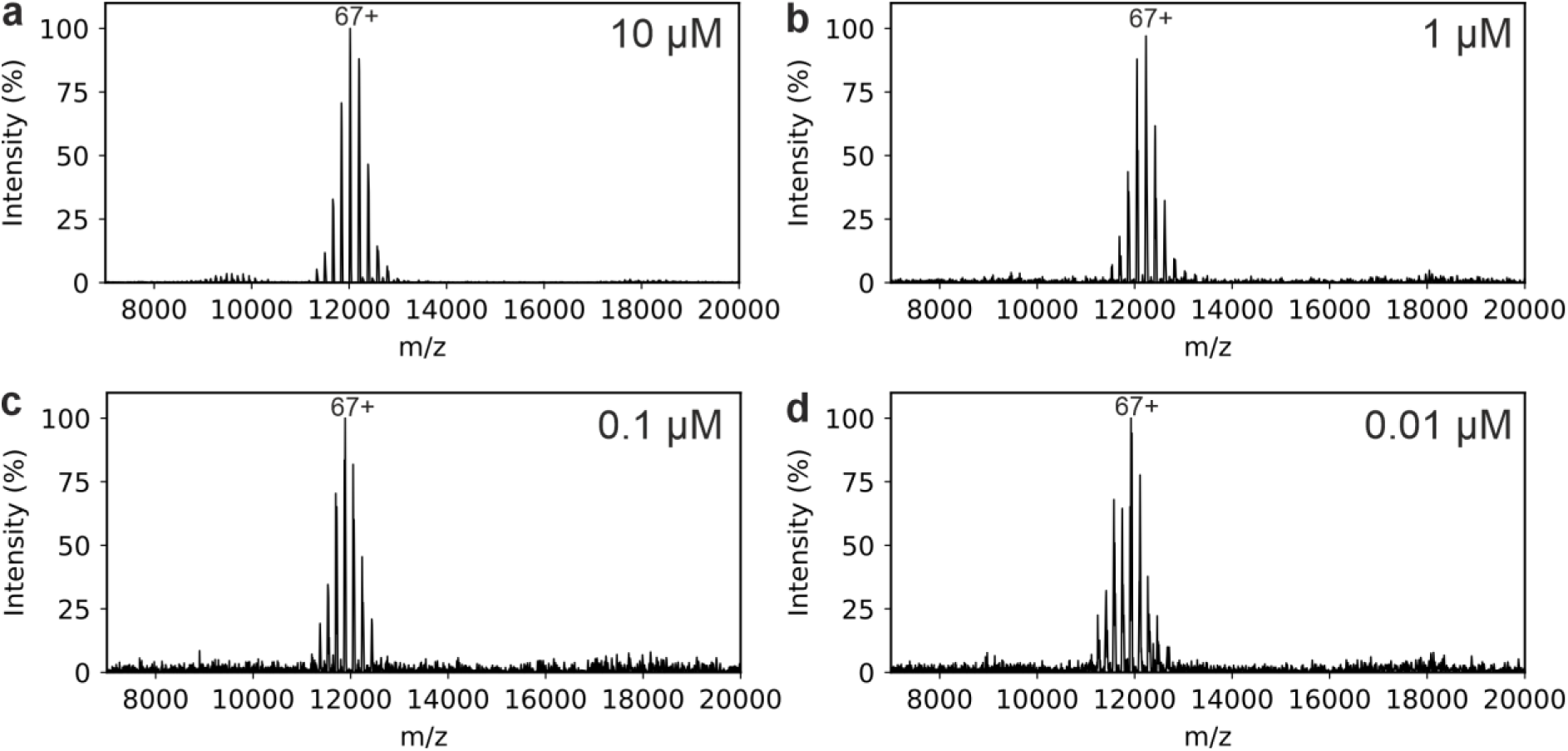
GroEL concentration series using nanopipette emitters. Native mass spectra of GroEL at protein concentrations of (a) 10 μM, (b) 1 μM, (c) 0.1 μM, (d) 0.01 μM. All spectra were acquired in 200 mM ammonium acetate (pH 6.6-6.8) on a UHMR Orbitrap.

Together with our transferrin, GDH and β-Gal datasets in Figure 4 and Figure 5, respectively, these results demonstrate that nanopipette nESI emitters extend routine, high-quality nMS into higher MW protein complexes/assemblies, retaining noncovalent interactions at sub-micromolar protein concentrations.

### Nanopipettes improve MDH dimer isolation

To test portability to a third MS platform, we analysed malate dehydrogenase (MDH; dimer ∼70 kDa) on an Orbitrap Ascend Tribrid instrument. With standard 1-2 µm emitters under typical native tuning conditions using an in-source collision induced dissociation (IS-CID) voltage of 200 V, the MDH native mass spectrum showed overlapping subunit/dimer charge state distributions, reflecting collisional activation and dimer dissociation in the gas phase as a result of the elevated IS-CID conditions needed to remove retained solvent (Figure 7a)^61,62^. Reducing the IS-CID below 200 V to reduce dimer dissociation was not possible whilst also maintaining a well-resolved mass spectrum. In comparison, nanopipette emitters at 1.0 kV capillary voltage and IS-CID 50 V enabled stable electrospray at markedly lower capillary potentials and reduced IS-CID, resulting in resolvable monomeric (blue) and dimeric (red) charge state envelopes that likely reflects a proportion of monomeric MDH in solution (Figure 7b) rather than the in-source monomer ejection observed with standard emitters (Figure 7a). Reducing the capillary voltage further to 0.4 kV increased the abundance of the MDH dimer relatively compared to the monomer (Figure 7c).

**Figure 7.**
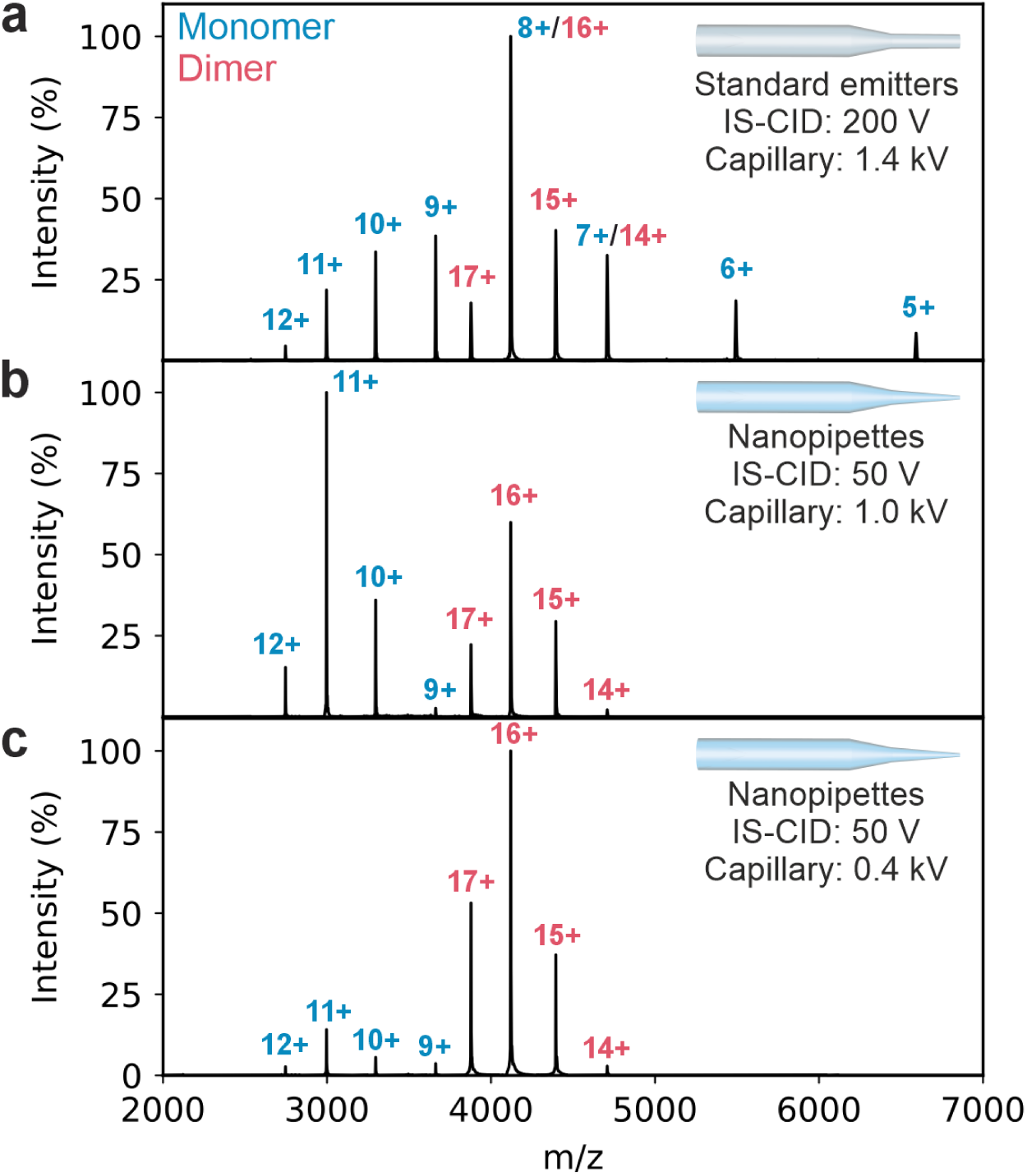
Nanopipette emitters lower required activation and preserve the native MDH dimer. (a) Mass spectrum of MDH acquired using a standard emitter (capillary 1.4 kV, IS-CID 200 V) has overlapping monomer (blue) and dimer (red) charge state distributions consistent with dimer dissociation in the gas phase. (b) Analysis of MDH using nanopipette emitters enables a well resolved mass spectrum using less activating source voltages (1.0 kV capillary, IS-CID 50 V), resulting in the detection of native-like monomer (blue) and dimer (red) charge state distributions. (c) Mass spectrum of MDH acquired using nanopipette emitters at lower source voltages than in (b) (0.4 kV capillary, IS-CID 50 V). Spectra were acquired in 150 mM ammonium acetate, pH 6.6-6.8.

### Nanopipette-enabled native MS of α-synuclein on the Bruker timsTOF Pro 2

To assess the transferability of nanopipette nESI to trapped ion mobility platforms, αS in 50 mM in ammonium acetate was analysed on a Bruker timsTOF Pro 2. Using nanopipette emitters (around 20 nm I.D.), αS produced charge-state-resolved native mass spectra with clear differences in spectral quality and adducting when compared to commercial online nESI emitters (20 μm I.D., Bruker). With both emitters, αS displayed a well-defined charge-state distribution consistent with monomeric protein (Figure 8a). However, using the standard commercial online nESI emitters increased salt adduction and peak broadening were observed (Figure 8b). Additionally, the charge state distribution appears to shift when comparing the MS spectra presented in Figure 8a, due to the lower capillary voltage applied in the case of nanopipettes, which reveals a more-native like charge state distribution whereby αS populates lower charge states (8+ - 6+; more compact conformations) more abundantly. Dimer peaks also appear to be present at higher intensities in Figure 8b, likely due to the lower capillary voltage applied, preserving the noncovalent assembly.

**Figure 8.**
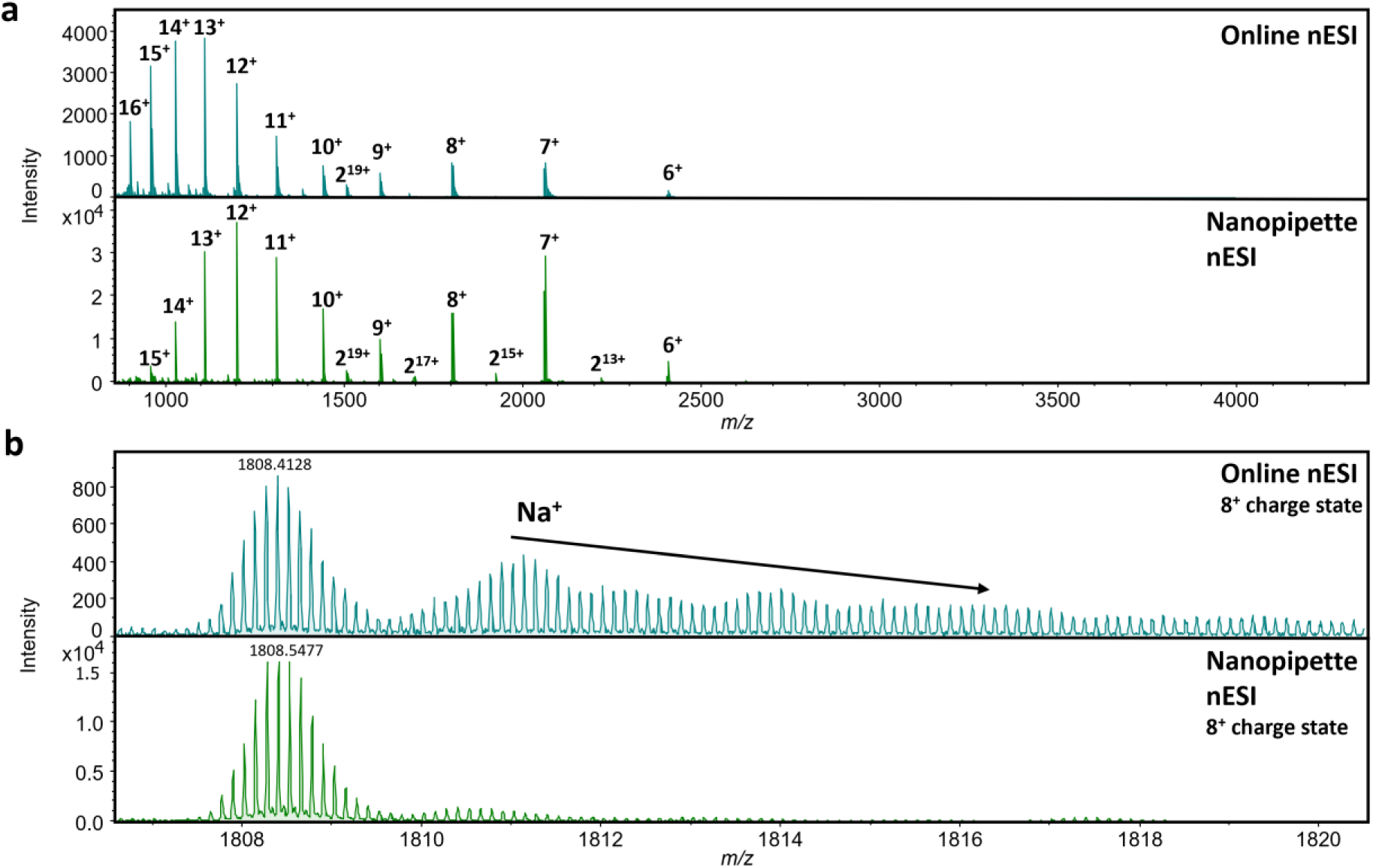
Native MS analysis of α-synuclein using nanopipette nESI on the Bruker timsTOF Pro 2. (a) Representative native mass spectrum of αS acquired using online nESI (blue, top spectrum) compared to nanopipette emitters (green, bottom spectrum). Charge states are denoted above spectral peaks. Peaks corresponding to αS dimers are labelled with a 2, with the charge state in superscript. (b) Zoom-in of the 8+ charge state, demonstrating that nanopipette nESI emitters improve desalting (green, bottom spectrum) compared to online nESI (blue, top spectrum) where Na^+^ adducts are observed.

The timsTOF platform also enabled ion mobility separation of αS conformational populations. The extracted mobility profile for the 8+ charge state revealed multiple partially resolved conformational families, consistent with the broad ensemble shown previously for the intrinsically disordered protein αS^44,45,63^. Figure S9 reveals high resolution native mass spectra where dimer populations can be observed for a particular conformational family within the 8+ charge state via the resolved isotopic distribution. These results demonstrate that nanopipette nESI can be implemented on the timsTOF Pro 2 for native analysis of intrinsically disordered proteins and provides a route to compare both mass spectral quality and ion mobility-resolved conformational heterogeneity directly from biologically relevant buffers.

### Complex matrices: bacterial lysates

We next investigated whether submicron nanopipette emitters enable native MS directly from crude mixtures. nMS has previously been applied directly to clarified bacterial and eukaryotic cell lysates to read out intact mass, oligomeric state, cofactors, and ligand binding without prior purification, enabling rapid characterisation of overexpressed targets directly from crude matrices^64,65^. Methods have since expanded towards exploration of drug-binding in lysates and to workflow variants such as online buffer exchange for lysates and even native top-down MS from tissue lysates, underscoring the feasibility of complex-matrix nMS^66,67^.

Here, *E. coli* lysates were prepared either co-expressing N-terminally acetylated α-synuclein (αSNTA) or a PET-28a vehicle control. Cells were harvested by centrifugation, resuspended in 100 mM ammonium acetate and lysed by sonication. Cell lysates were then centrifuged to remove particulates, and the supernatant was diluted 1:1 with 100 mM ammonium acetate, as outlined in Figure S10, before analysis on a Synapt G2-Si instrument. Using nanopipette nESI emitters, the lysates from cells expressing αSNTA produced a resolved αSNTA monomer charge state distribution spanning charge states 6+ to 14+ (Figure 9a), with dominant, resolvable peaks and minimal nonspecific adducts despite the matrix background (Figure S11). In contrast, the pET-28a vehicle control lysate showed no corresponding peaks under otherwise identical instrument parameters, with lowly abundant peaks likely arising from endogenous *E. coli* proteins (Figure 9b). Spectra acquired using standard borosilicate emitters are shown in Figure S12. An observable post-translational modification can be identified to αSNTA charge states which corresponds to a D-gluconyl modification which occurs in *E. coli* over time^68^.

**Figure 9.**
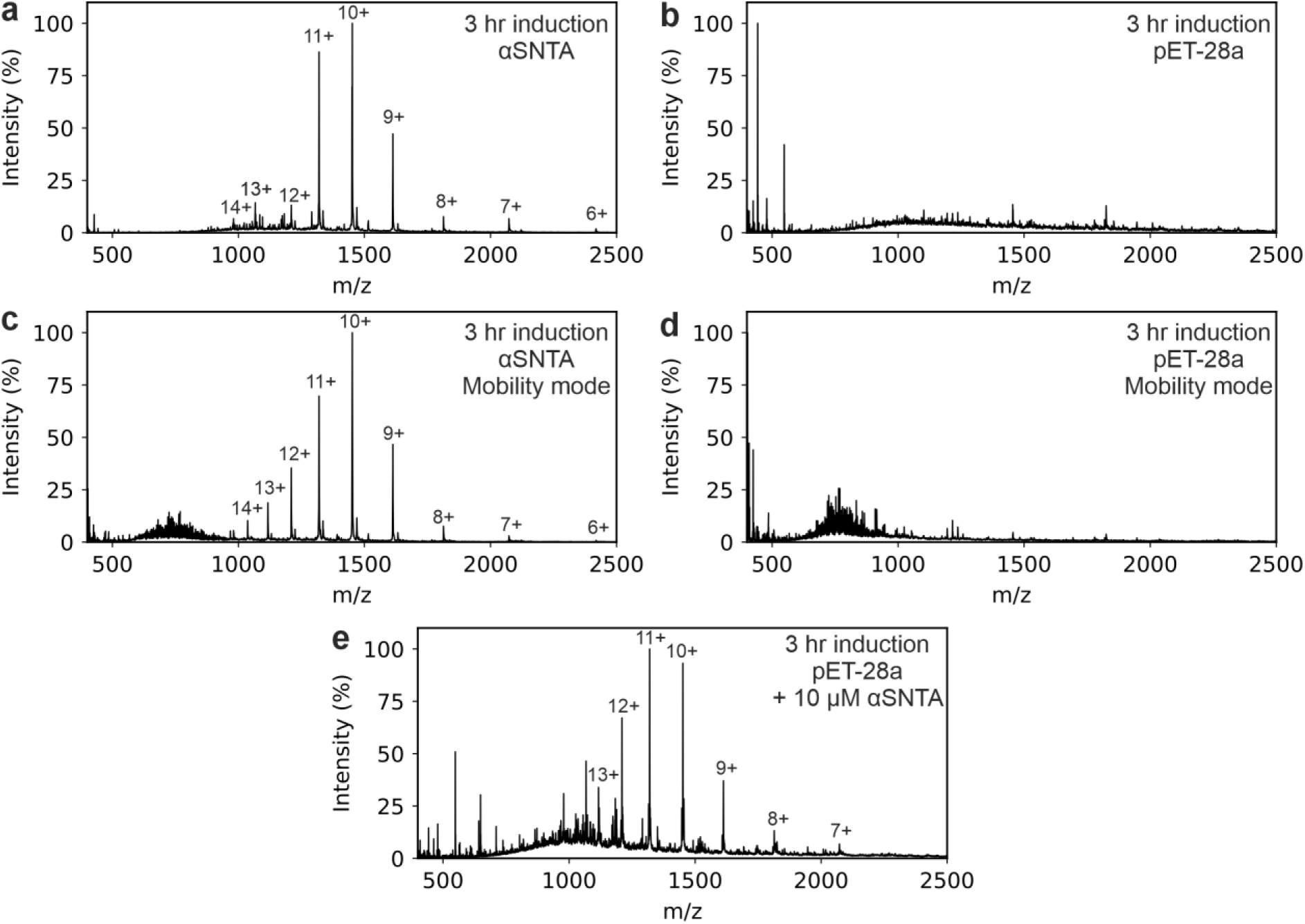
Native MS of N-terminally acetylated α-synuclein directly from clarified lysates with submicron nanopipette nESI emitters. (a) Induced lysate expressing N-acetylated α-syn (αSNTA) with a resolved monomer charge state distribution (6+ to 14+). (b) Empty-vector vehicle control (pET-28a) under matched conditions. Background signal is from endogenous E. coli proteins. (c) Induced αSNTA lysate acquired in mobility mode. (d) Empty-vector vehicle control acquired in mobility mode. (e) PET-28a vehicle control lysate spiked with 10 μM α-SNTA. All data were collected on a Synapt G2-Si in 100 mM ammonium acetate, pH 6.6-6.8 using sub-50 nm quartz nanopipette emitters.

Acquiring in ion-mobility mode identifies an identical αSNTA charge state distribution (6+ to 14+; Figure 9c) and characteristic arrival time distribution with four evident conformational families, consistent with previously published calibrated collision cross section values (Figure S13)^44,45,63^ which were not observed in the pET-28a vehicle control (Figure 9d). Finally, spiking αSNTA (10 μM spike in concentration) directly into the PET-28a vehicle control lysate yielded a charge state distribution corresponding to αSNTA (Figure 9e). These data show that nanopipette nESI maintains proteoform-level information directly for proteins that are overexpressed in crude cell lysates, while ion mobility provides an orthogonal filter for identification and characterization, extending nanopipette enabled native MS beyond purified protein preparations.

## Conclusion

Submicron nanopipette nESI emitters provide a simple, portable route to highly resolvable, high signal-to-noise nMS across instruments and protein mass ranges. By benchmarking proteins from 17 to 800 kDa on a Waters Corporation Synapt G2Si, ThermoFisher Scientific Orbitrap UHMR, Orbitrap Ascend and Bruker timsTOF pro 2, we show that <50 nm quartz nanopipette emitters deliver native mass spectra of proteins at markedly lower capillary voltages and activation conditions, with systematically fewer nonspecific salt adduct peaks and higher signal to noise than conventional 1-2 μm borosilicate emitters. These demonstrations extend from small, standard proteins (αS, myoglobin, CAH II, ovalbumin) to larger noncovalent assemblies (MDH, transferrin, GDH, β-Gal, GroEL), and translate across instrument vendors indicating that performance reflects nanopipette-mediated desolvation from finer droplets rather than instrument-specific optimization. In practical contexts, nanopipettes preserve noncovalent cofactor interactions and protein assemblies and enable direct analysis of clarified lysates, preserving proteoform information. Together, these data establish nanopipettes as a robust front end for routine and challenging nMS measurements.

Our results provide operating guidance (tip size and instrument voltage conditions) and motivate systematic applications such as challenging buffer compositions and matrices and coupling to orthogonal readouts (ion mobility/collision induced unfolding, top-down fragmentation approaches and direct mass technology). We anticipate nanopipette nESI emitters will lower barriers to native MS in physiological buffers and complex samples across laboratories and applications.

## Author contributions

E.J.B fabricated quartz nanopipette emitters. E.J.B, E.J.O, Z.J.H, R.S, K.H, I.A, M.K, and L.W performed experiments and analysis. E.J.B, P.A, A.M.R, P.S, K.F, F.S, R.R.O.L, A.N.C, and J.A.L. developed the ideas and supervised the project. All authors contributed to the preparation of the manuscript.

## Acknowledgements

A.N.C. and E.J.B. acknowledge support through a Sir Henry Dale Fellowship jointly funded by Wellcome and the Royal Society (220628/Z/20/Z). EJB acknowledges support from the European Union’s Horizon Europe research and innovation programme through a Marie Skłodowska-Curie Actions Global Postdoctoral Fellowship (Project NANOSPEC, Grant Agreement No. 101274120. P.A., and A.N.C. acknowledge funding from the BBSRC (BB/X003086/1) and F.S., P.A. from the MRC (MR/W031515/1). P.A. acknowledges funding from the BBSRC (UKRI1930). J.A.L. acknowledges support from US National Institutes of Health (NIH) (R35GM145286). E.J.O. acknowledges NIH-NIGMS (T32GM145388) for support. A.M.R. gratefully acknowledges funding from the research program VICI with project number VI.C.192.024 and Aspasia (015.015.009) from the Dutch Research Council (NWO). P.S. acknowledge support from UC Riverside Startup Fund and NIH (R00AI183290). We thank Dr. Alexander Kulak (University of Leeds) for the SEM imaging of the emitters.

