## Supplemental Information for "Cross-Platform Assessment of Sub-50 nm Nanopipette Emitters for Native Electrospray Ionization Mass Spectrometry"

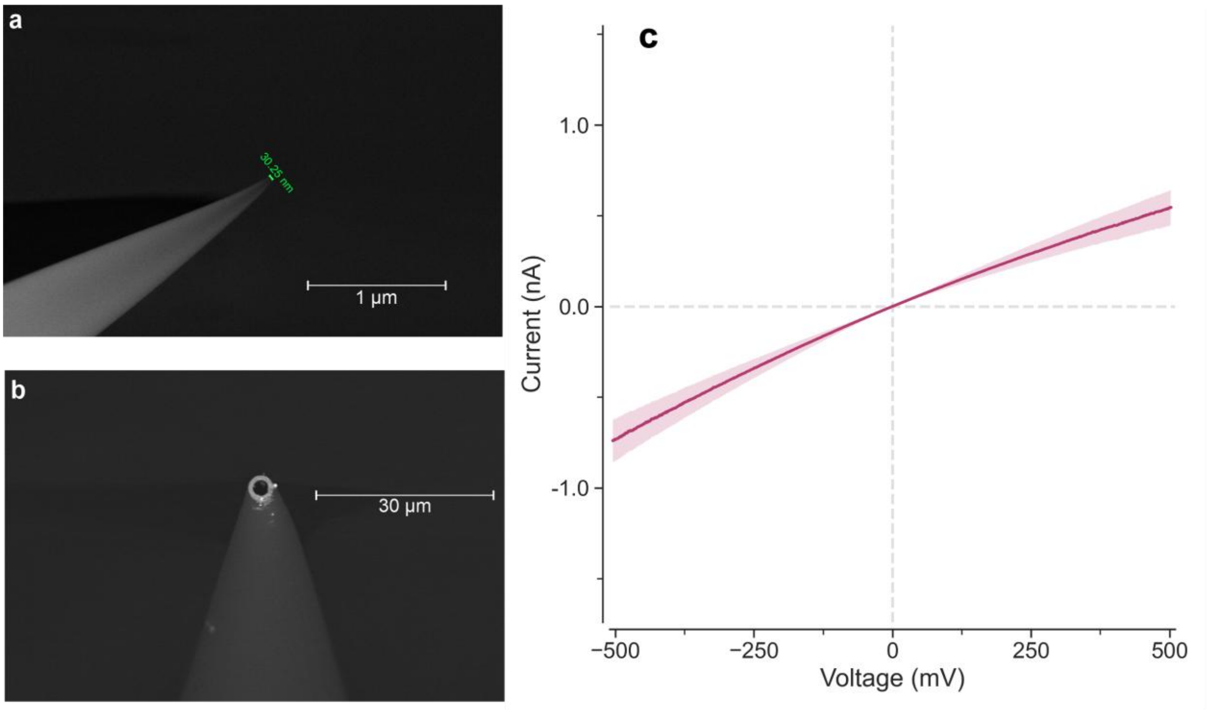


***Figure S1 Representative scanning electron microscopy micrographs of emitters and electrical characterization of the nanopipette emitters.*** *(a) Nanopipette quartz emitters, inner diameter: ~ 30 nm. (b) Standard borosilicate emitters, inner diameter: ~ 2 µm. (c) i-V curves of 6 quartz nanopipettes recorded in 0.1M KCl. The shaded area represents the standard error of the mean and the solid line is the average i-V curve of the nanopipettes.*

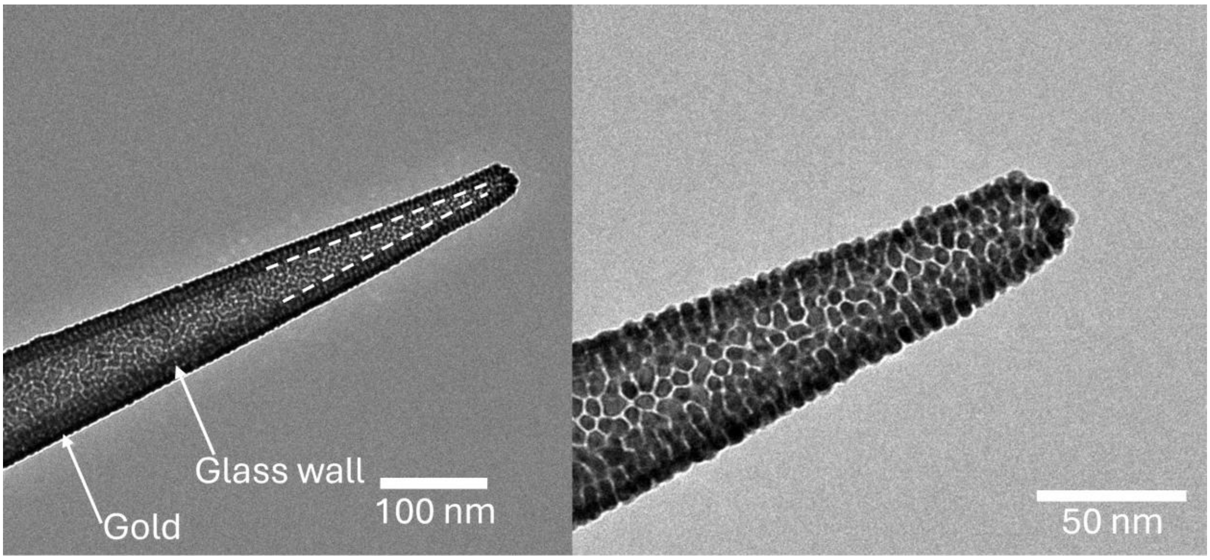


***Figure S2 Representative transition electron microscopy micrograph of coated nanopipette emitters.*** Nanopipettes (I.D. ~20 nm) were coated with a thin titanium adhesion layer followed by gold deposition.

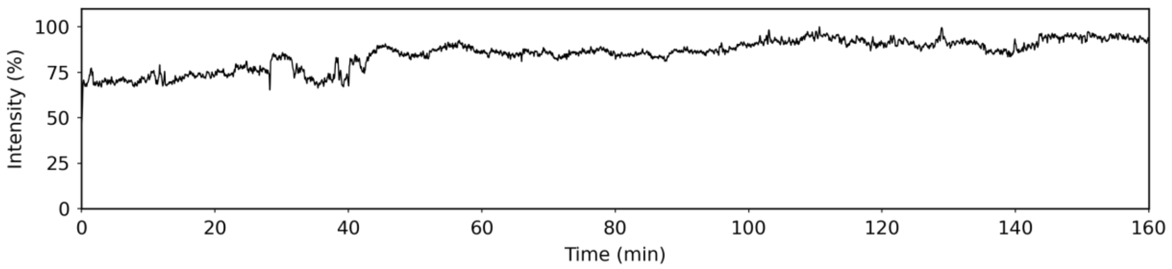


***Figure S3 CAH II stable electrospray chromatogram.*** *Electrospray signal for CAH II over 160 minutes acquired using a Waters Corporation Synapt G2Si.*

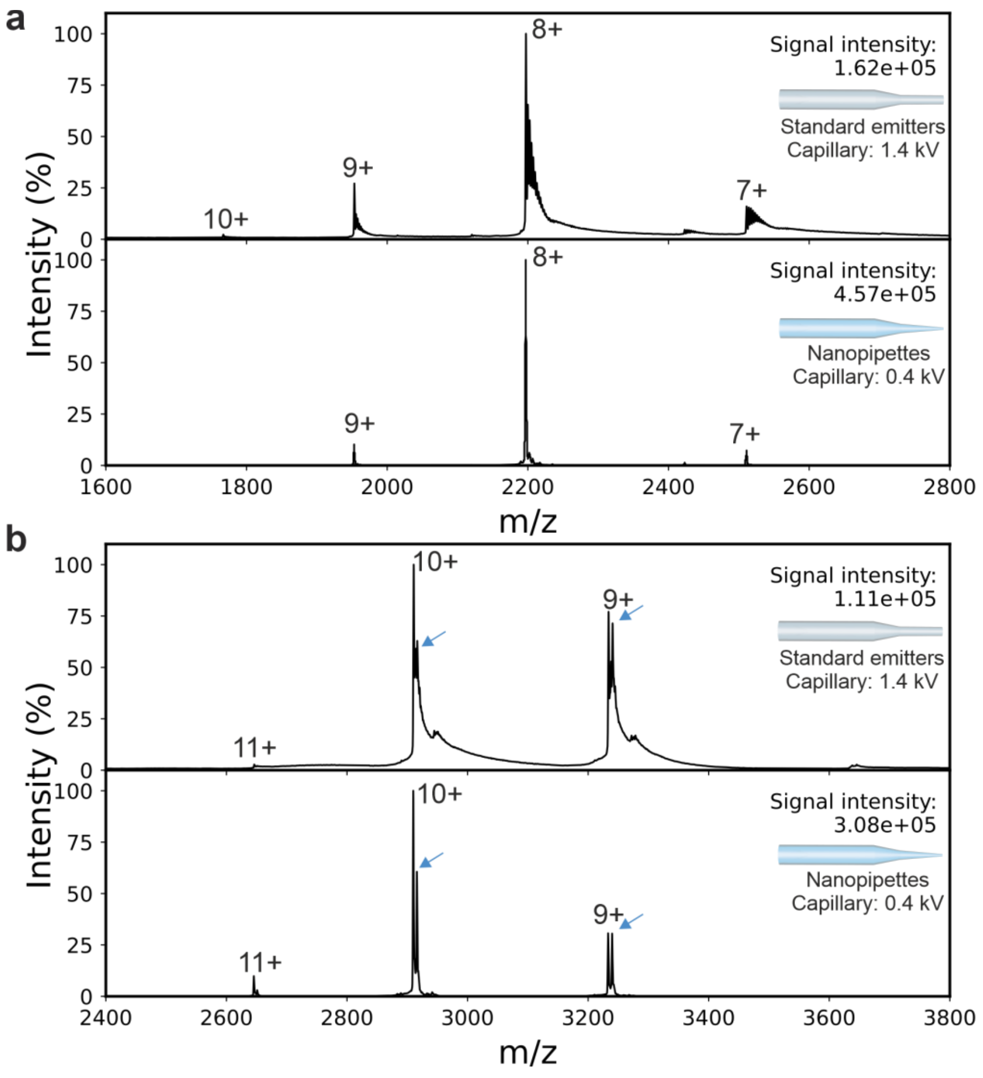


***Figure S4 Native mass spectra of myoglobin and CAH II using nanopipettes and standard ESI emitters.*** *Native mass spectra acquired using nanopipettes (above) and standard emitters (below) for myoglobin (a) and CAH II (b). Blue arrows indicate the* $Zn^{2+}$ *bound metalloproteoform. Total ion signal intensities and capillary voltages are shown on the right-hand side.*

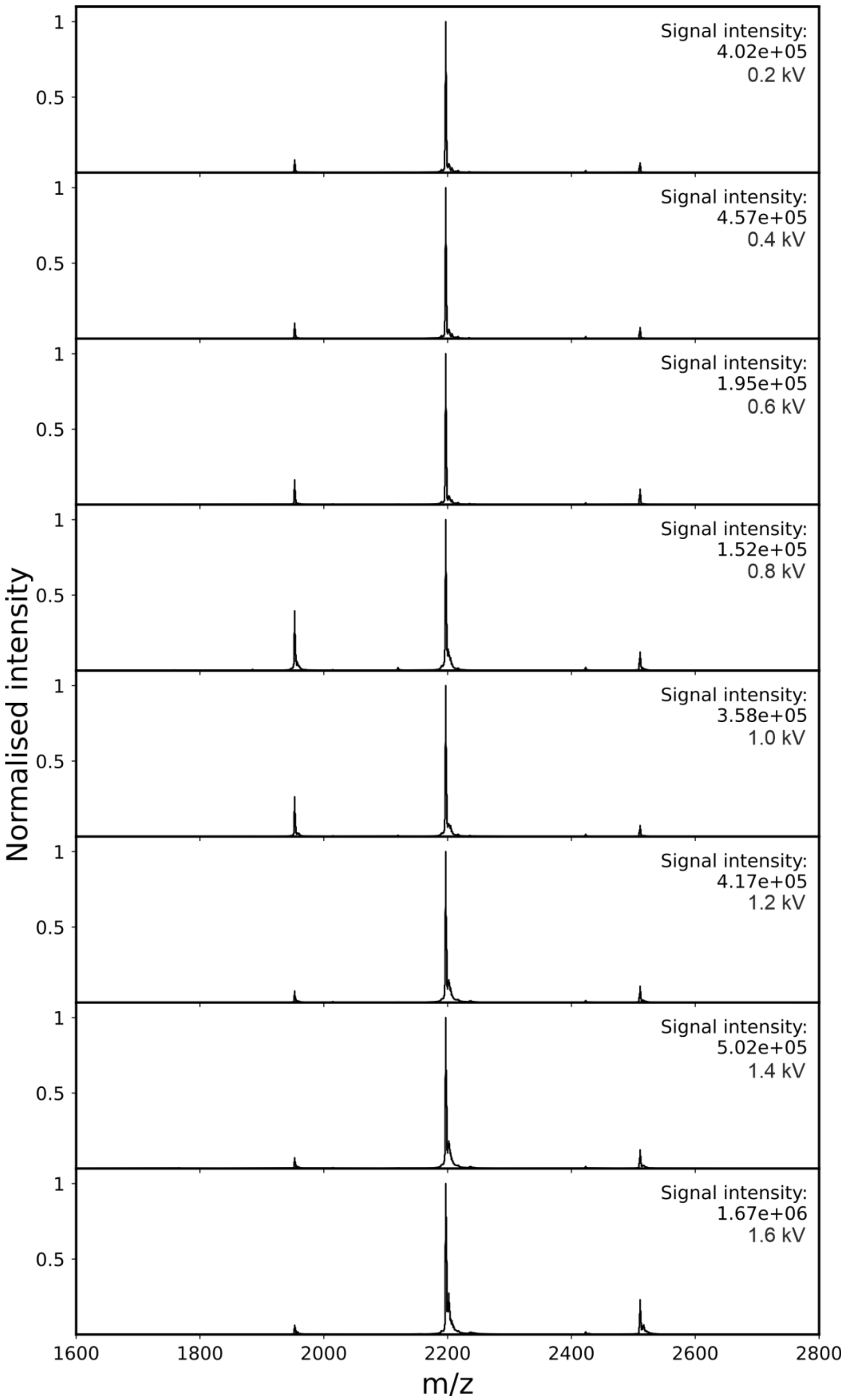


***Figure S5 Analysis of capillary voltage on spectral quality using nanopipette emitters for analysis of myoglobin.*** *Representative native mass spectra of myoglobin acquired using nanopipettes ranging from 0.2-1.6 kV capillary voltage from top to bottom. Data were acquired as two-minute acquisitions using a Waters Corporation Synapt G2Si.*

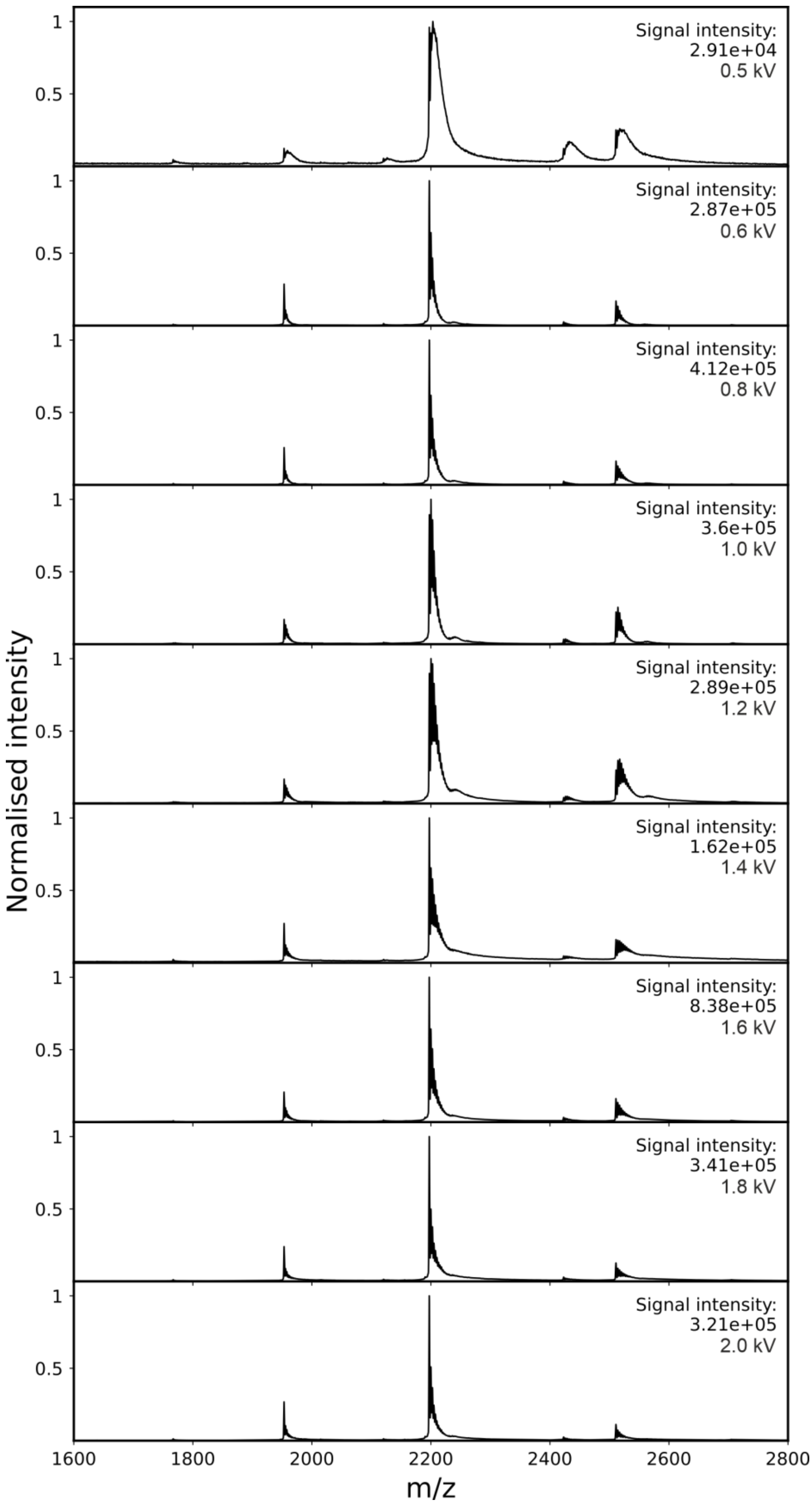


***Figure S6 Analysis of capillary voltage on spectral quality using standard emitters for analysis of myoglobin.*** *Representative native mass spectra of myoglobin acquired using nanopipettes ranging from 0.5-2.0 kV capillary voltage from top to bottom. 0.5 kV was the lowest voltage that signal could be observed. Data were acquired as two minute acquisitions using a Waters Corporation Synapt G2Si.*

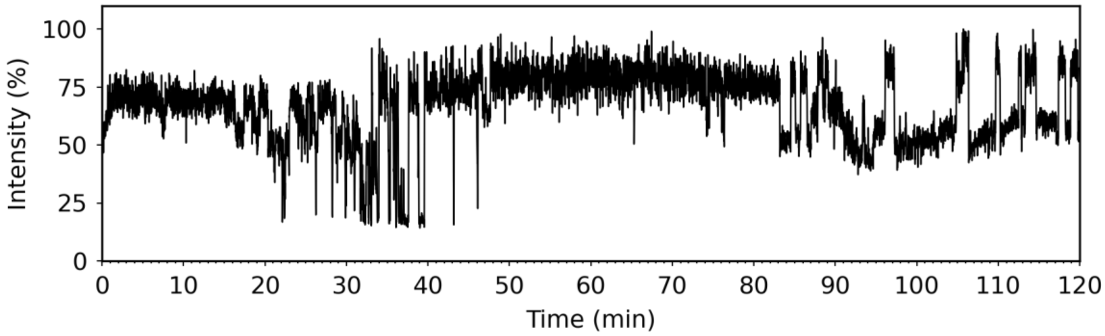


***Figure S7 β-Galactosidase stable electrospray chromatogram.*** *Electrospray signal for β-Galactosidase over 120 minutes acquired using a ThermoFisher Scientific Q Exactive UHMR Orbitrap.*

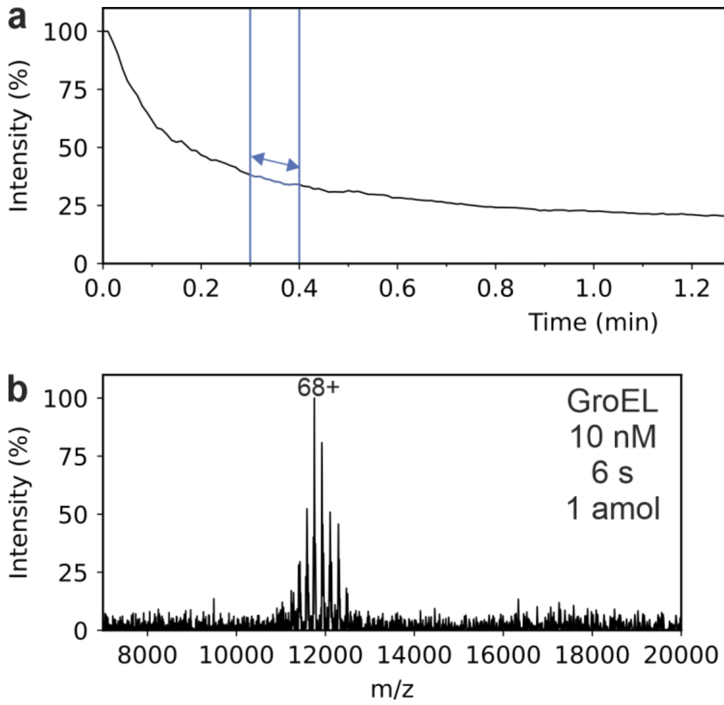


***Figure S8 Acquisition of 1 amol of GroEL.*** *(a) Total ion chromatogram for 10 nM GroEL acquisition, the blue arrow indicates a six second window combined to generate the (b) native mass spectrum.*

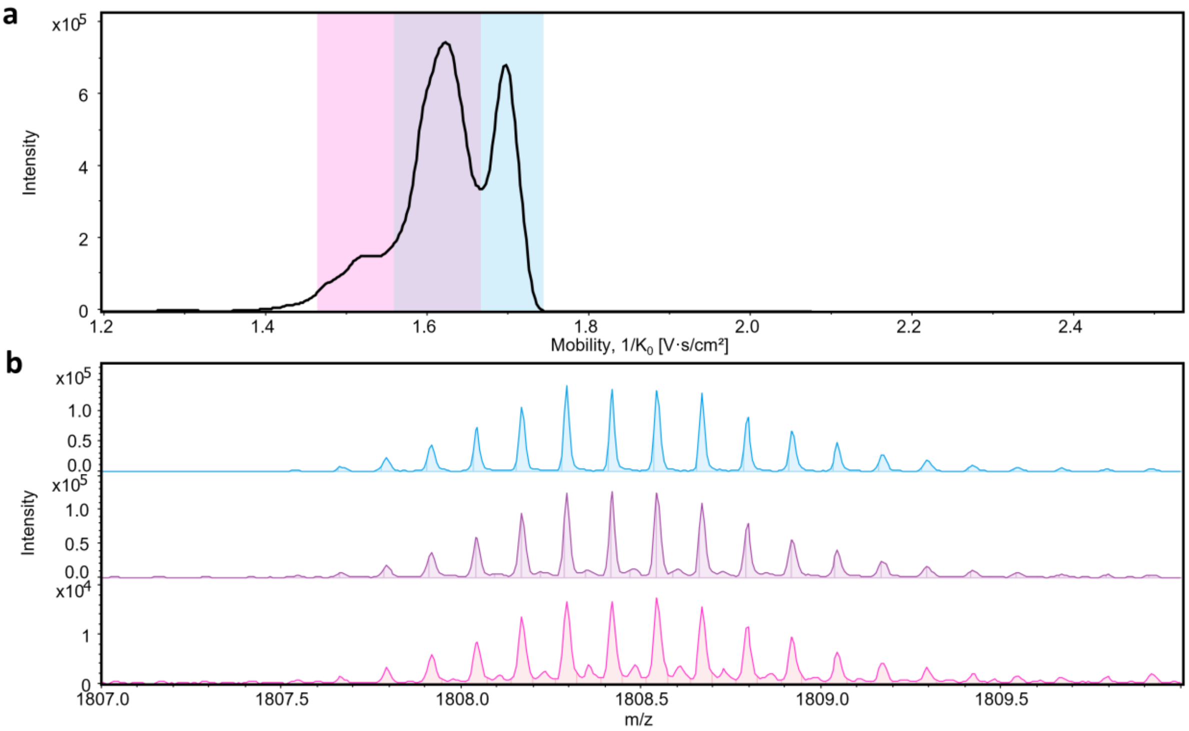


***Figure S9. Ion mobility-resolved conformational analysis of αS on the Bruker timsTOF Pro 2.*** *(a) The extracted ion mobility profile of the 8+ charge state of αS and (b) representative native mass spectra of αS from specific conformational families acquired using a nanopipette nESI emitter. The mobility distribution reveals multiple partially resolved conformational families, consistent with the heterogeneous ensemble of intrinsically disordered αS*[44,45,59]*. A dimer population can be observed from the most compact conformational family (pink) which is resolved using isotopic distribution in the high resolution mass spectrum (pink, b)*

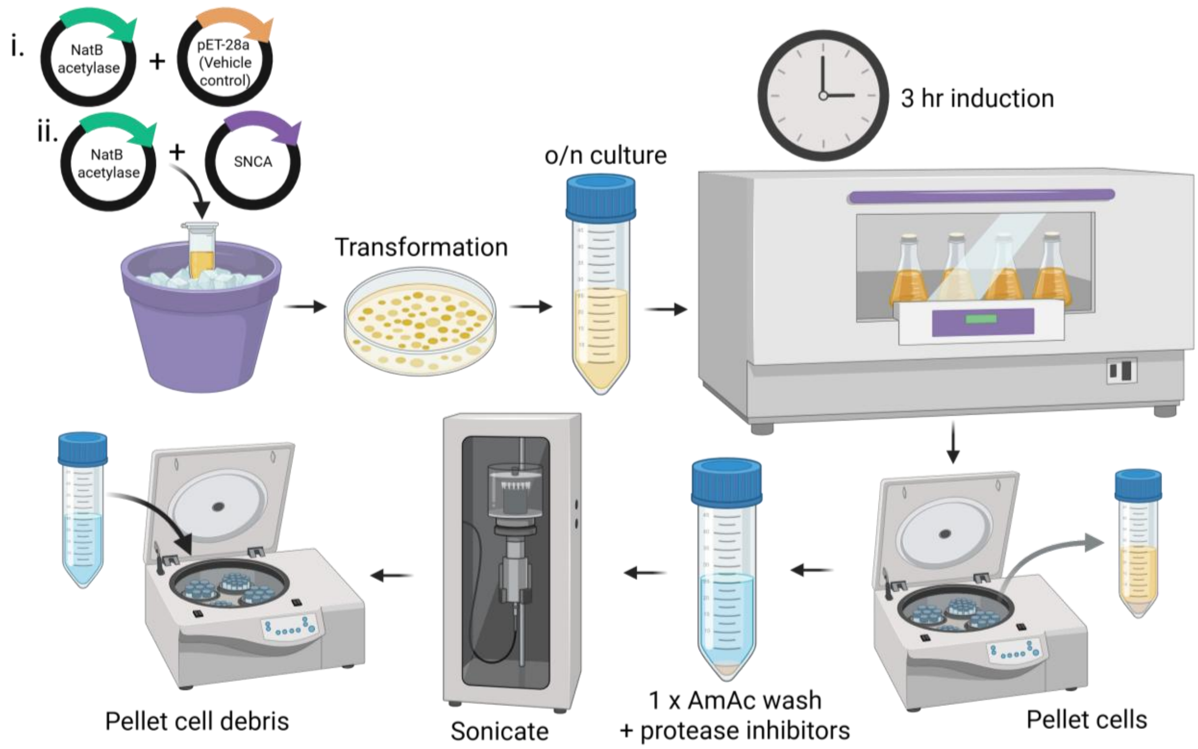


***Figure S10 Workflow for producing clarified bacterial lysates for nMS.*** *Illustrated steps used to generate lysates for direct analysis by nanopipette nESI (see Methods for exact conditions). Briefly the steps include transformation of competent cells with the expression construct (α-synNTA or pET-28a vehicle control) and overnight growth of colonies. The overnight culture was used to inoculate 50 mL LB cultures and protein expression was initiated by IPTG induction for 3 hours. Cells were harvested by centrifugation and resuspended in 100 mM ammonium acetate before pelleting again. Cell lysis was performed by sonication with the addition of protease inhibitors. Cell debris were removed through centrifugation.*

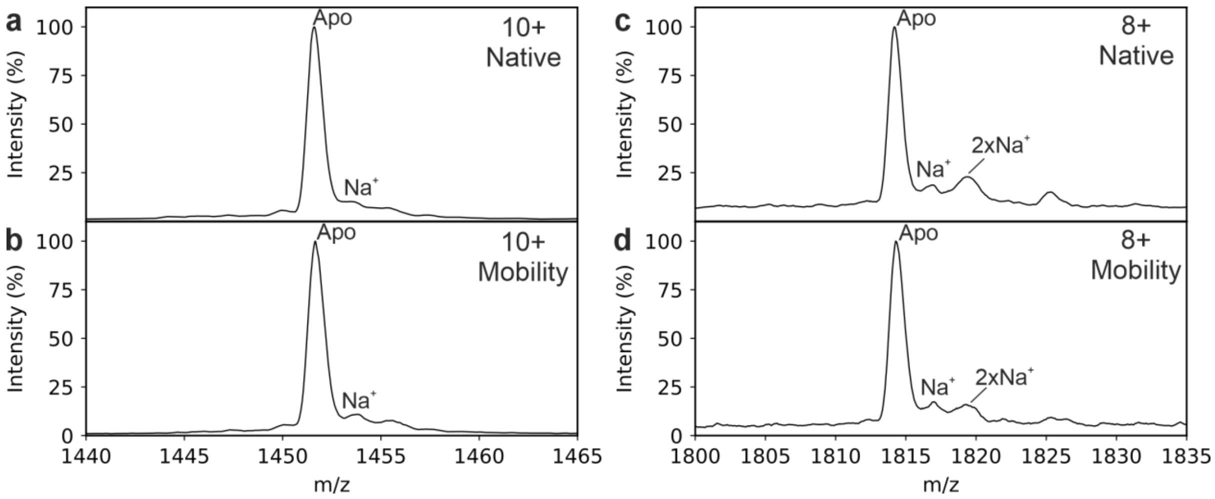


***Figure S11 Zoomed in native mass spectra from bacterial lysates recombinantly expressing N-terminally acetylated α-synuclein.*** *Induced lysate expressing N-acetylated α-syn (αSNTA) 10+ charge state in MS mode (a) and mobility mode (b) and for the 8+ charge state in (c) MS mode and (d) mobility mode, measured using a Waters Corporation Synapt G2-Si instrument.*

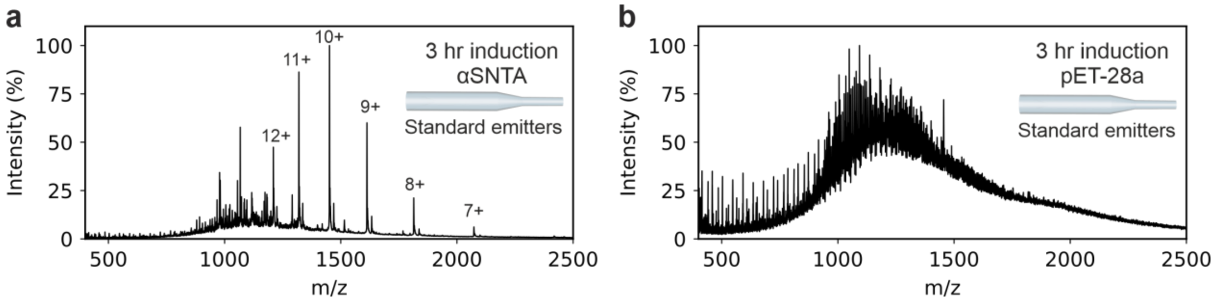


***Figure S12 nMS analysis of lysate from standard emitters.*** *(a) Induced lysate expressing N-acetylated α-syn (αSNTA) with a resolved monomer charge state distribution (6+ to 12+). (b) Empty-vector vehicle control (pET-28a) under matched conditions.*

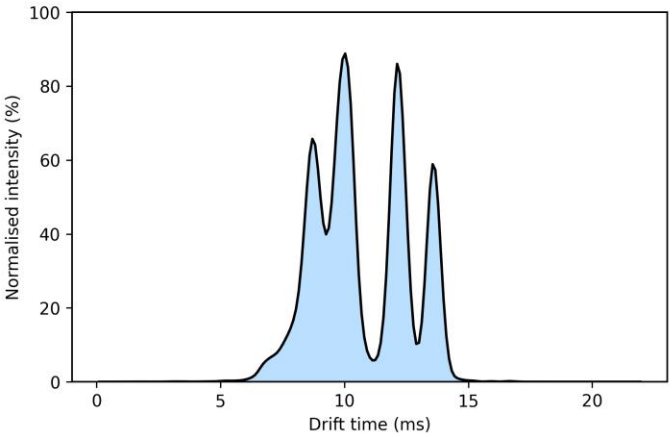


***Figure S13 Arrival time distribution of the 8+ charge state of recombinantly expressed N-terminally acetylated α-synuclein directly from bacterial lysates.*** *Arrival time distribution of the 8+ charge sate of αSNTA measured in ms reflects the previously observed distribution of the 8+ charge state from calibrated ion mobility measurements*[41,42,60].
